# Extracellular vesicles produced by pathogenic *E. coli* expressing *hlyF* compromise epithelial barrier function

**DOI:** 10.64898/2026.11.25.769135

**Authors:** Marie Pénary, Christelle Marrauld, Eric Oswald, Priscilla Branchu

**Affiliations:** Université de Toulouse, INSERM, INRAE, ENVT, IRSD, Toulouse, France; CHU Toulouse, Hôpital Purpan, Service de bactériologie-hygiène, 31024 Toulouse, France

## Abstract

The intestinal epithelium constitutes a critical barrier against toxins, microbiota, and pathogens, and its integrity is essential for maintaining host homeostasis. Selected pathogens or pathobionts, under specific conditions, have the capacity to disrupt barrier functionDespite its close genetic similarity to the adherent-invasive *Escherichia coli* (AIEC) strain LF82, NRG857c has a greater capacity to colonize the intestine and induce colitis in conventional wild-type mice. The NRG857c plasmid differs from that of LF82 and encodes HlyF, an enzyme known to modulate the formation and properties of bacterial extracellular vesicles (bEVs). Here, we demonstrated that bEVs derived from the AIEC strain NRG857c disrupted the intestinal barrier integrity with no detectable increase in LDH release after 5 h of exposure, in an *in vitro* Caco-2 epithelial model. We showed that this effect is specifically dependent on HlyF, a cytoplasmic enzyme from the short-chain dehydrogenase-reductase superfamily. We identified that EVs produced by BL21 (DE3) expressing the active form of *hlyF* significantly decreased transepithelial electrical resistance (TEER) in Caco-2 cells epithelium. This reduction in TEER was associated with tight junction disorganization and an increased paracellular permeability. In contrast, EVs derived from bacteria expressing a mutant in the catalytic site of *hlyF* did not affect TEER, tight junction architecture, or permeability, remaining comparable to untreated controls. Altogether, our results demonstrated that *hlyF*-dependent EVs produced by *E. coli* compromise intestinal epithelial barrier integrity. These findings highlight a novel mechanism by which pathobiont-derived vesicles contribute to barrier dysfunction and may promote intestinal inflammation under pathological conditions.

**Author summary:** The intestinal epithelial cells form a barrier that limits the crossing of substances from the lumen of the gut into surrounding tissues. This barrier is altered and become more permeable in Crohn’s disease patients. Adherent and invasive *Escherichia coli* are associated with Crohn’s disease. We asked whether nanoparticles released by these bacteria, called vesicles, could alter the barrier. We found that vesicles from one such strain made a layer of human intestinal cells grown in the laboratory more permeable. In addition, vesicles produced by bacteria with an active form of an enzyme called HlyF disrupted the structures that seal neighboring cells together. Vesicles produced by bacteria with an inactive form of HlyF left the barrier intact. Our findings showed that bacterial vesicles can weaken the intestinal barrier on their own. Our study reveals a previously unrecognized way in which bacterial vesicles weaken the intestinal barrier through a process that depends on the activity of the bacterial enzyme HlyF.

## Introduction

Adherent-invasive *Escherichia coli* (AIEC) have been implicated in intestinal epithelial barrier dysfunction. Increased epithelial permeability may facilitate bacterial access to the lamina propria, sustaining inflammation and creating conditions that favor AIEC persistence [1–3]. Conversely, AIEC infection has been shown to alter epithelial architecture, modulate tight junction proteins and impair epithelial integrity [4,5]. Altogether, these observations suggest that barrier dysfunction is both a consequence and a driver of AIEC colonization. However, the molecular mechanisms responsible for these alterations remain incompletely understood and are unlikely to rely solely on direct AIEC–epithelium interactions.

Most studies investigating AIEC pathogenicity have focused on bacterial adhesion, epithelial invasion and intracellular survival [3,6–14]. Nevertheless, these contact-dependent mechanisms cannot fully explain how relatively localized bacterial populations induce widespread epithelial dysfunction within the intestinal mucosa. This observation raises the possibility that diffusible bacterial factors capable of acting remotely from the bacterial cell contribute to epithelial barrier impairment. Identifying these mechanisms is therefore essential to better understand how AIEC remodel the intestinal microenvironment and promote chronic inflammation.

An emerging concept in bacterial pathogenesis is that host manipulation is not exclusively mediated by direct bacteria-host cell interactions. Gram-negative bacteria constitutively release bacterial extracellular vesicles (bEVs), that are nanosized lipid bilayer structures originating from the outer membrane that function as highly efficient vehicles for the delivery of bacterial molecules [15–19]. First considered as by-products of bacterial growth, bEVs are now recognized as sophisticated secretion platforms capable of transporting proteins, lipids, lipopolysaccharides, peptidoglycan fragments, metabolites and nucleic acids to host cells without the need of direct contact [15–21].

The nanoscale size of bEVs enables them to diffuse through biological fluids and mucus layers, potentially cross epithelial barriers and interact with multiple cell types within host tissues [17–19,22]. Consequently, bEVs extend the sphere of bacterial influence far beyond the immediate site of colonization, allowing pathogens and pathobionts to remotely modulate epithelial physiology, innate immunity and inflammatory responses [17,19,22]. Depending on their molecular composition, bacterial vesicles could activate pattern-recognition receptors, deliver toxins, interfere with intracellular trafficking pathways and reshape host signaling networks [15,23–25]. They have therefore emerged as major mediators of long-distance host–microbe communication.

Increasing evidences indicate that vesicles produced by enteric bacteria participate in the regulation of intestinal homeostasis. Several studies have shown that bEVs influence epithelial signaling, cytokine production and junctional organization, ultimately affecting epithelial permeability [24,26,27]. However, these biological activities vary considerably between bacterial species and strains, suggesting that their function may result from diverse mechanisms governing vesicle biogenesis and cargo composition [18,20,21]. The bacterial determinants responsible for generating bioactive vesicle populations remain poorly understood.

Interestingly, recent studies have begun to implicate bEVs produced by AIEC in host-pathogen interactions, including inflammatory responses and altered expression of junction-associated proteins in intestinal epithelial cells [5]. However, whether these vesicles directly contribute to epithelial barrier dysfunction and which bacterial factors determine their pathogenic activity remain largely unknown. Addressing these questions is essential for understanding how AIEC remotely remodel the intestinal environment and promote chronic intestinal inflammation.

The biological activity of bEVs is intimately linked to their molecular composition, yet the bacterial determinants governing vesicle functionality remain largely unknown [18,20,21]. Among extraintestinal pathogenic *Escherichia coli* (ExPEC), we previously identified HlyF as a virulence factor promoting the production of bEVs with peculiar properties against eucaryotic cells [28]. Despite its historical annotation as a putative hemolysin, HlyF is a cytoplasmic enzyme belonging to the short-chain dehydrogenase/reductase (SDR) superfamily that promotes the production of bEVs displaying pro-inflammatory properties [28,29]. Using complementary cellular and animal models, we previously demonstrated that HlyF-dependent vesicles inhibit autophagic flux, potentiate non-canonical inflammasome activation and exacerbate host cell cytotoxicity, revealing that HlyF profoundly remodels host–pathogen interactions through modulation of bEVs properties rather than by acting as a conventional secreted virulence factor [28,29].

Interestingly, the AIEC strain NRG857c harbors an *hlyF*-containing virulence plasmid that is absent from the prototype strain LF82 [28,30]. Moreover, NRG857c displays an enhanced ability to colonize the intestine and promote experimental colitis compare to LF82 [31]. These observations led us to hypothesize that HlyF-dependent bEVs may contribute to this increased pathogenic potential by targeting the intestinal epithelial barrier. However, whether HlyF-dependent bEVs directly impair epithelial barrier integrity has never been investigated.

In the present study, we demonstrate that HlyF-dependent bEVs produced by *E. coli* disrupt intestinal epithelial barrier function by decreasing transepithelial electrical resistance (TEER), increasing paracellular permeability and inducing tight junction disorganization. Our findings identify HlyF as a key bacterial determinant controlling the pathogenic activity of bEVs and uncover a previously unrecognized mechanism by which AIEC-derived bEVs may contribute to epithelial dysfunction and intestinal inflammation independently of direct bacterial-epithelial contact.

## Results

### Deletion of *hlyF* abolishes the barrier-disrupting activity of bEVs from AIEC strain NRG857c

We differentiated Caco-2 cells in epithelium on insert for three weeks and verified the barrier function by measuring the transepithelial electrical resistance (TEER) before treatment. We harvested and purified bEVs produced by the AIEC strain NRG857c, the mutant Δ*hlyF* and the complemented mutant Δ*hlyF* (pAGO15). Epithelia were treated with bEVs from the different strains and the TEER was measured at different time point post-treatment up to 4h30. The TEER dropped specifically when epithelia were treated with bEVs from the NG857c wild-type or the complemented mutant Δ*hlyF* (pAGO15) (**Figure *1***A). The TEER of the epithelia treated with bEVs from the mutant Δ*hlyF* was similar to the TEER measured in the untreated epithelia. These results showed that a treatment of Caco-2 cells epithelia with bEVs from the AIEC NRG857c dropped the TEER, and that this drop is specifically due to the presence of *hlyF* in the genome of NRG857c. After 5 h of exposure to wild-type NRG857c bEVs, LDH release from Caco-2 monolayers was not significantly different from untreated controls, suggesting that the decrease in TEER occurred without detectable cytotoxicity at this time point. However, we observed a significantly higher LDH release when the Caco-2 monolayers were treated with bEVs from the complemented mutant Δ*hlyF* (pAGO15), suggesting a dose-effect on the cytotoxicity.

**Figure 1:**
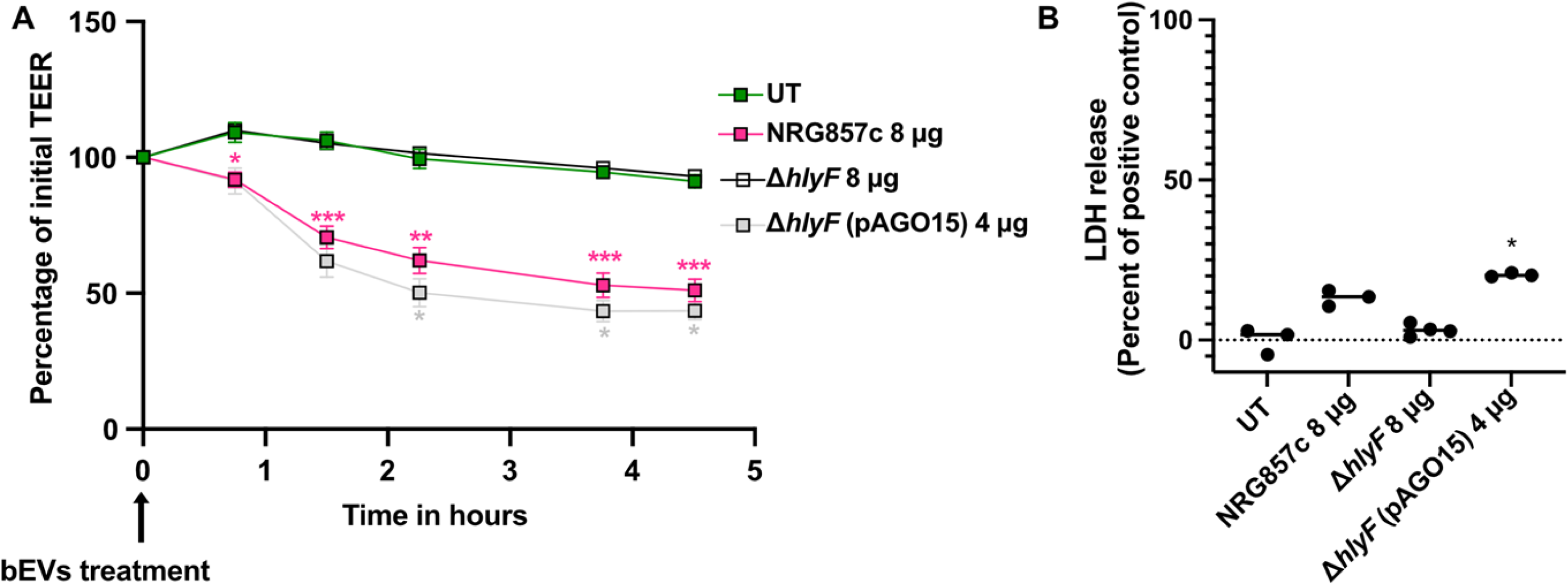
Effects of NRG857c-derived bEVs on TEER and LDH release in Caco-2 cells monolayers. Caco-2 cells monolayers were exposed to bEVs derived from wild-type NRG857c, the mutant Δ*hlyF* or the complemented mutant Δ*hlyF* (pAGO15). A-Transepithelial electrical resistance (TEER) across the monolayers. B-Lactate dehydrogenase (LDH) release from the monolayers. bEVs expressed in μg of proteins. Statistical significance was assessed using two-way repeated-measures ANOVA followed by Dunnett’s multiple comparisons test versus the untreated condition (UT).

### Mutation of the HlyF catalytic site abolishes the ability of bEVs to disrupt the intestinal epithelial barrier

To avoid side-effects that could be due to other virulence genes present in the AIEC strain NRG857c, we introduced *hlyF* or its mutant in the catalytic site Y163F-K167A (HlyF_Y163F-K167A_) in the laboratory *E. coli* strain BL21(DE3). We purified bEVs from each derived strain (named bEVs-HlyF and bEVs-HlyF_Y163F-K167A_, respectively) and treated Caco-2 epithelial monolayers. We observed that the TEER of Caco-2 cells epithelia dropped specifically when they were treated with 12,5 μg or 25 μg of bEVs-HlyF (**Figure 2**A). Conversely, the TEER of Caco-2 cells epithelia were similar to untreated epithelia when they were treated with bEVs from BL21(DE3) expressing the mutant in the catalytic site or 5 μg of bEVs produced by the BL21 (DE3) expressing the functional version of *hlyF*. These results showed that the drop of the TEER observed when the epithelia are treated with bEVs is specific of the modification induced by the activity of HlyF. To identify if the drop of the TEER is due to a mortality of Caco-2, we estimated the viability of Caco-2 monolayers after 5 hours treatment by measuring the LDH release in the supernatant in the different conditions. LDH release remained comparable to that of untreated cells under all conditions, suggesting that the decrease in TEER was unlikely to result from bEVs-induced cytotoxicity.

**Figure 2:**
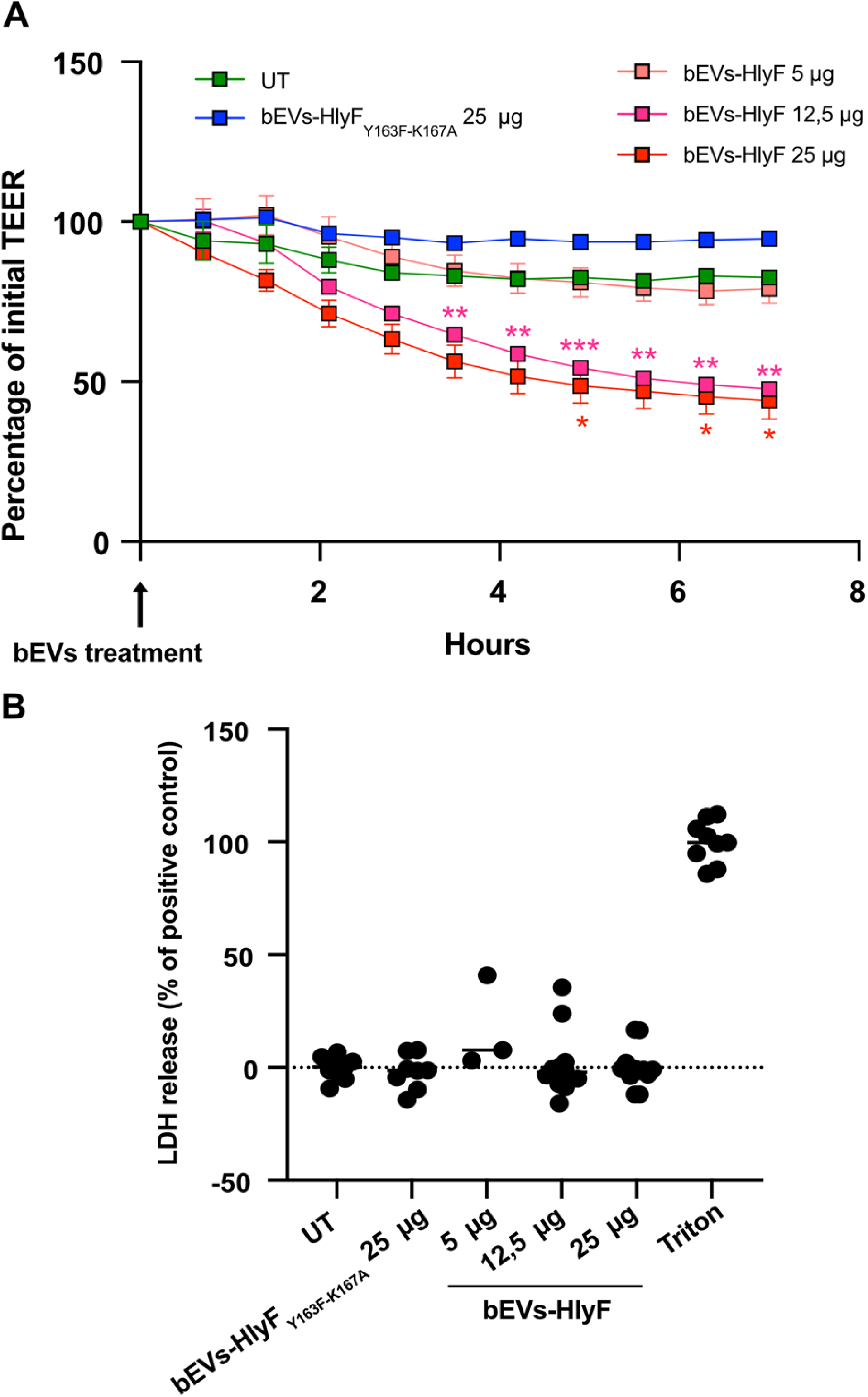
bEVs produced by a non-pathogenic strain disrupt the epithelial integrity without inducing mortality. A-TEER measured after treatment with different concentrations of bEVs produced by the non-pathogenic strain BL21(DE3) expressing the functional version of *hlyF* or the mutant in the catalytic site Y163F-K167A. B-Release of LDH in the supernatant of Caco-2 monolayers in the different conditions after 5 hours treatments. Results are expressed in percentage of positive control (cell lyzed with triton).

### The disruption of the intestinal epithelium induced by bEVs from an *E. coli* producing HlyF increases the paracellular permeability

A decrease in TEER is often associated with an increase of paracellular permeability. To identify if bEVs-HlyF have the capacity to increase paracellular permeability, we assessed the crossing of the fluoresceine sodium sulfate (FSS) and FITC-Dextran through Caco-2 monolayers treated or not with bEVs-HlyF or the control bEVs-HlyF_Y163F-K167A_. We didn’t observe differences in permeability with untreated Caco-2 monolayers when they were treated with 25 μg of control bEVs-HlyF_Y163F-K167A_ or 5 μg of bEVs-HlyF (**Figure 3**). However, with a treatment of 12,5 μg bEVs, permeability was significantly increased for sodium fluorescein (FSS, approximatively 376 Da), whereas no significant increase was detected for 4-kDa FITC-dextran. Notably, permeability was significantly increased for both tracers when Caco-2 monolayers were treated with 25 μg of bEVs-HlyF, indicating a dose-dependent impairment of epithelial barrier permeability.

**Figure 3:**
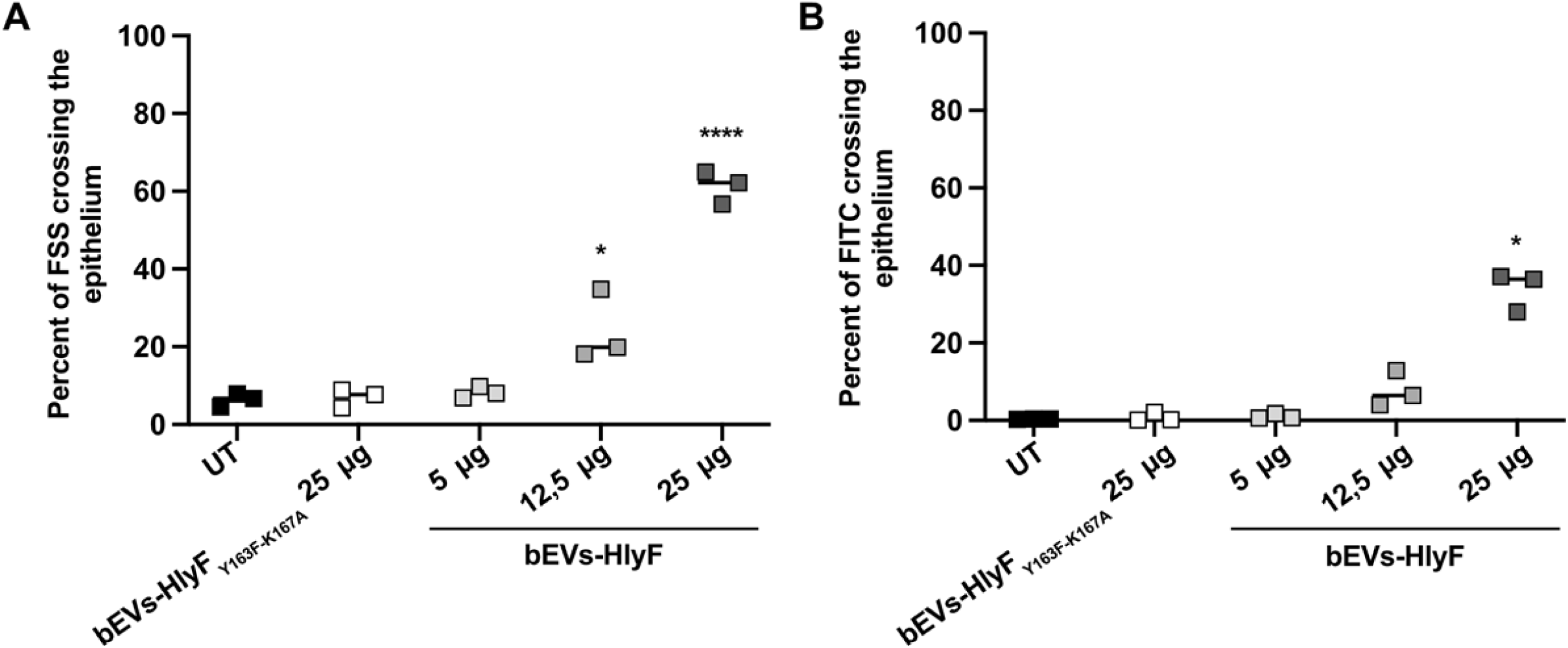
Paracellular permeability to sodium fluorescein and 4-kDa FITC-Dextran following treatment with bEVs. Caco-2 monolayers were treated with bEVs-HlyF or bEVs-HlyFY163F-K167A or left untreated (UT). FSS or FITC-dextran was added to the apical compartment 24h before bEVs treatment and its passage into the basolateral compartment was assessed 24h after treatment. Results are expressed as the percentage of FSS or FITC-dextran crossing the membrane relative to cell-free inserts. Data are from three biological replicates.

**Figure 4:**
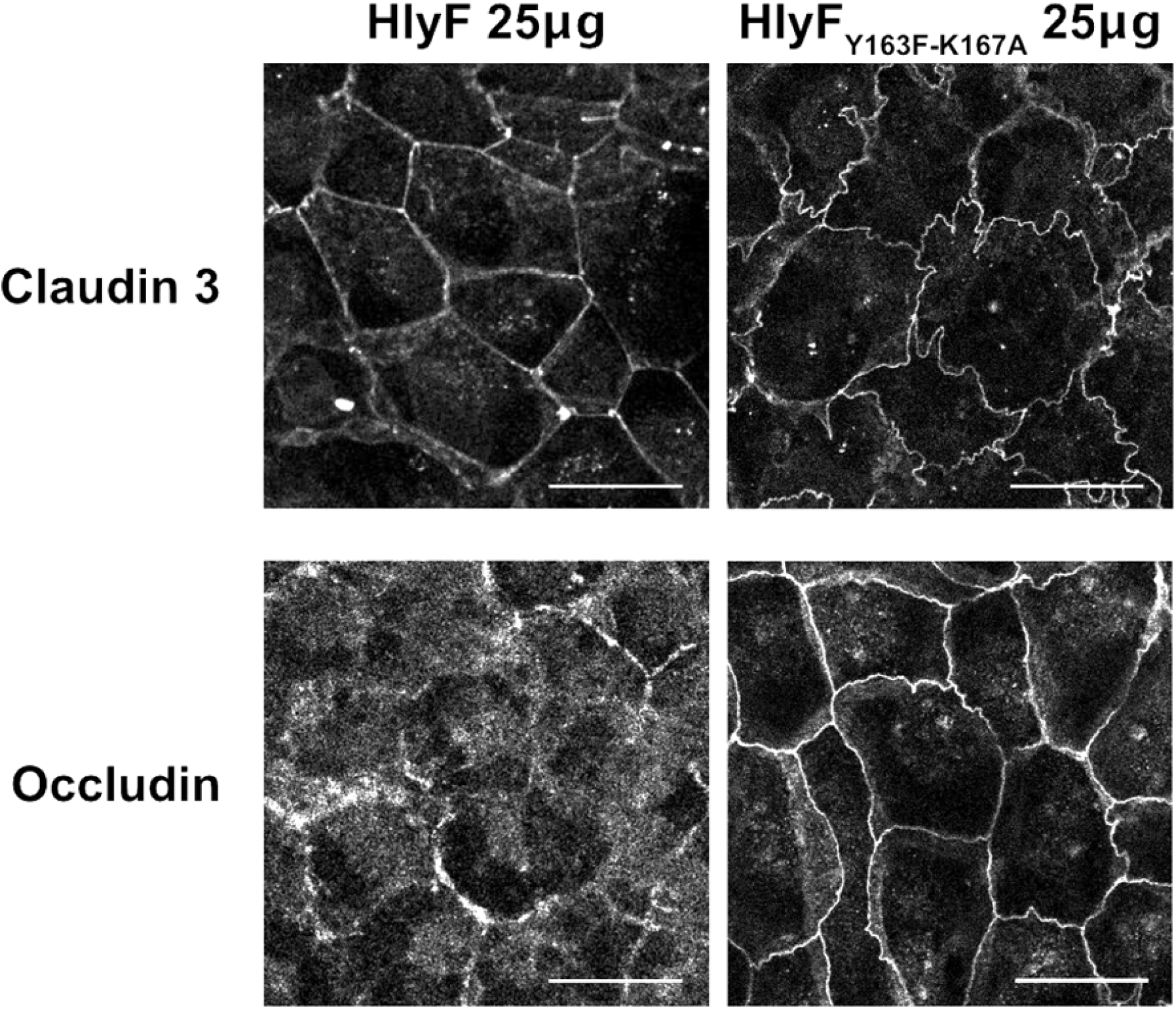
Immunostaining of Tight Junctions in Caco-2 cells monolayers treated with bEVs. Caco-2 cells monolayers were treated with 25 μg of bEVs derived from the strain producing functional HlyF enzyme or mutated if the catalytic site HlyF_Y163F-K167A_ followed by an immunostaining of the tight junction proteins Claudin 3 or Occludin. Images were acquired by confocal laser scanning microscopy and analyzed with Fiji. Scale bar 20 μm.

### bEVs from an *E. coli* producing HlyF disrupt the tight junctions of the intestinal epithelium

We showed that the decrease in TEER is not due to a mortality and is associated with an increase of paracellular permeability. We previously demonstrated that bEVs-HlyF specifically block the autophagy flux in HeLa cells. Previous studies demonstrated that a functional autophagy help maintaining the barrier function by reducing the degradation of tight junction proteins such as occluding [32,33]. To test if the increase of paracellular permeability observed specifically when Caco-2 monolayers were treated with bEVs-HlyF is due to a disorganization of tight junctions, we performed immunostaining against Occludin and Claudin 3 after a treatment of the Caco-2 monolayers with bEVs-HlyF or bEVs-HlyF_Y163F-K167A_. Confocal microscopy revealed distinct patterns of tight-junction proteins between the two conditions. Claudin 3 staining displayed differences in the organization of cell boundaries following treatment with bEVs-HlyF compared with bEVs-HlyF compared with bEVs-HlyF_Y163F-K167A_. More strikingly, occluding displayed a more diffuse distribution following bEVs-HlyF treatment, whereas a sharply defined and continuous junctional staining was observed in monolayers treated with bEVs-HlyF_Y163F-K167A_. These observations indicate that the catalytic activity of HlyF is associated with modifications in tight-junction protein organization.

## Discussion

AIEC have been associated with the ileal mucosa of patients with Crohn’s disease [6], and the reference strain LF82 has been shown to disrupt apical junctional complexes and impair barrier function in polarized epithelial monolayers [4]. Although NRG857c and LF82 share similar chromosomal backgrounds, they differ in their virulence plasmid content, with NRG857c hosting a plasmid that harbours *hlyF* [34], a gene absent from LF82. This distinction is relevant to the interpretation of their vesicular effects: the epithelial responses reported for LF82-derived bEVs occur independently of HlyF, whereas the decrease in TEER induced by NRG857c-derived bEVs required *hlyF* under our experimental conditions. Our findings therefore identify a contribution of HlyF within the diversity of bacterial determinants underlying AIEC-epithelial interactions.

Previous work on LF82-derived bEVs has primarily examined their contribution to bacterial invasion and epithelial responses. Rolhion and colleagues demonstrated that these vesicles participate in epithelial interactions promoting invasion of the bacteria [35]. Nadalian and colleagues subsequently reported inflammatory responses and changes in the expression of junction-associated genes in Caco-2 cells exposed to LF82-derived bEVs [5]. Their main analyses used 10 µg/mL OMVs for 24 h in cultures at 80-90% confluence and assessed junction-associated responses at the transcript level. Our study complements these observations by examining the functional integrity of an epithelial monolayer through TEER measurement, tracer passage, together with junctional protein organization. This distinction matters because changes in junction-associated gene expression and changes in barrier function describe different levels of the epithelial responses. The combined findings support a role for AIEC-derived vesicles in epithelial perturbation, while showing that the contribution of individual bacterial determinants must be considered within each strain background.

The biological relevance of this phenotype is particularly apparent in the context of Crohn’s disease. Increased intestinal permeability may precede clinical diagnosis. Indeed, a prospective study of asymptomatic first-degree relatives showed that impaired barrier function was associated with subsequent disease development, including when measured more than three years before diagnosis [36]. Experimental evidence also indicates that increased tight junction permeability can promote mucosal immune activation and contribute to experimental colitis [37]. These observations place epithelial barrier dysfunction at the interface between microbial exposure and host susceptibility, where it may contribute to disease development and progression rather than solely reflect established tissue injury.

Within this framework, the ability of NRG857c-derived bEVs to impair barrier function provides a potential link between an AIEC-associated bacterial determinant and increased mucosal exposure to luminal products. LDH release following exposure to wild-type NRG857c-derived bEVs was comparable to that of untreated controls at 5 h. The altered organization of occludin and claudin-3 is consistent with a disturbance of intercellular junctions that could weaken the functional separation between the intestinal lumen and underlying tissues. In a susceptible host, such a change could increase immune exposure to microbial products and contribute to the persistence or amplification of mucosal inflammation. Isolated bEVs were sufficient to elicit this epithelial response in our model. Vesicle-mediated effects may therefore complement bacterial adhesion and invasion in shaping the interactions of AIEC with the intestinal mucosa.

This interpretation places HlyF within the bacterial processes that determine the biological activity of bEVs. Previous work established its contribution to bEVs biogenesis [28]; our findings connect its functionality to an epithelial barrier phenotype. Because the comparisons were performed at equivalent protein doses, they establish differences in the activity of the bEVs preparations under this normalization. How HlyF functionality translates into these differences, and which bEVs-associated components mediate the epithelial response, remain questions for understanding the link between bEVs production and host effects. The present results establish this link at the functional level without assigning HlyF-bEVs a direct action on epithelial junctions.

The significance of vesicle-associated barrier alterations may extend beyond the intestinal epithelium. Studies of *Porphyromonas gingivalis* bEVs have documented increased endothelial permeability and altered vascular junctions, as well as reduced barrier resistance and disrupted ZO-1 organization in human blood–brain barrier models [38,39]. These findings place bacterial vesicles within a broader framework of interactions with epithelial and endothelial barriers. Our results add to this framework by identifying HlyF functionality in the producing bacterium as a determinant of the intestinal barrier response. They raise the question of whether HlyF-associated bEVs effects are restricted to the intestinal epithelium or may also influence vascular and blood-brain barriers. Considering both the bacterial determinants of bEVs activity and the properties of the host barrier may therefore help explain how bEVs contribute to distinct outcomes across tissues.

## Materials and methods

### Bacteria strains, mutagenesis and growth conditions

The bacterial strains used in this study are listed in Table S1. Bacteria were routinely grown in lysogeny broth (LB, Lennox). For bEVs production, bacteria were grown in M63 minimal medium (Bio Basic) supplemented with 0.2% D-glucose (G8769, Sigma) and 0.1 mM MgCl₂ (83266, Sigma-Aldrich). When required, kanamycin sulfate (UK0015, Euromedex) was added at 50 mg/L.

The NRG857c Δ*hlyF* mutant was generated by λ Red-mediated homologous recombination using the method described by Datsenko and Wanner [40]. Briefly, an FRT-flanked kanamycin resistance cassette was amplified from pKD4 using primers carrying extensions homologous to the regions flanking *hlyF*. The resulting PCR product was introduced into NRG857c expressing the λ Red recombination system from pKD46, allowing replacement of *hlyF* with the resistance cassette. Recombinant clones were selected on kanamycin, and the deletion was verified by PCR. The kanamycin resistance cassette was subsequently excised using FLP recombinase expressed from pCP20.

### Purification of bacterial extracellular vesicles

Bacterial precultures were initiated from glycerol stocks stored at −80°C and grown overnight in LB supplemented with antibiotics when required. Overnight cultures were diluted 1:100 into supplemented M63 medium and incubated at 37°C for 8 h. Cultures were centrifuged at 6,000 rpm for 10 min, and the resulting supernatants were passed through 0.22-µm filters. Filtered supernatants were concentrated by tangential flow ultrafiltration using membranes with a molecular weight cutoff of 100 kDa (Vivaflow 200, VF20P4, Sartorius), followed by ultracentrifugation at 32,000 rpm for 3 h. The resulting bEVs pellets were resuspended in phosphate-buffered saline (PBS). Protein concentrations in the bEVs preparations were determined with the BCA protein assay kit using bovine serum albumin as a standard (DCTM Protein Assay Kit II, 5000112, Biorad).

### Cell cultures and bEVs treatment

Caco-2 cells (ATCC, HTB-37) were seeded onto Transwell inserts (Falcon, 353095, 0,4µm pore size, 0,3 cm^2^) at a density of 20,000 cells per insert in Dulbecco’s modified Eagle medium (DMEM) supplemented with 10% fetal bovine serum (FBS), 1% non-essential amino acids (MEM NEAA, 11140-035, Gibco) and 1% Penicilline/Streptomycin (P4333, Sigma). Cells were maintained at 37°C with 5% CO₂ concentration, and the medium in both the apical and basolateral compartments was replaced every other day. Monolayers were allowed to differentiate for 3 weeks, and their integrity was assessed by measuring transepithelial electrical resistance (TEER).

On the day before treatment, the medium in both compartments was replaced with DMEM supplemented with 5% FBS. Monolayers were subsequently exposed to bEVs in the apical compartment at the indicated doses and for the indicated durations. Doses were expressed as µg of bEVs-associated protein per insert.

### Measurement of transepithelial electrical resistance

TEER was measured using a cellZscope Plus system (Nano Analytics distribute by Protéigène). Values were normalized to the corresponding pretreatment value (T₀) and expressed as percentages using the following equation: Relative TEER (%) = (TEER at time t / TEER at T₀) × 100.

### Paracellular permeability assays

On the day before treatment, the culture medium was replaced with DMEM supplemented with 5% FBS and containing either FITC-dextran at 5 mg/mL (MW: 389,38) or FSS at 0,4 mg/mL (MW: 376,27) in the apical compartment. Differentiated Caco-2 monolayers were subsequently treated with bEVs at the indicated doses.

Tracer passage across the monolayers was assessed by measuring fluorescence in the basolateral compartment at 24h, using the SPARK plate reader (Tecan) with excitation/emission wavelengths of 485/530 nm, respectively, for FITC-dextran and FSS. Results were expressed as percentage of FITC-Dextran or FSS crossing the insert relative to cell-free inserts.

### Immunofluorescence and microscopy analysis

Caco-2 monolayers grown on Transwell inserts were washed three times with PBS and fixed either with 3.7% paraformaldehyde for 30 min at 4°C for occludin staining or with methanol for 1 min at −20°C for claudin-3 staining. Following three washes with PBS, cells were permeabilized with 0.5% Triton X-100 in PBS for 5 min at room temperature. Monolayers were then washed three times with PBS containing 0.1% Tween, with the final wash lasting 10 min. Nonspecific binding was blocked with 10% goat serum in PBS containing 0.1% Tween for 30 min at room temperature.

Following three washes with PBS, monolayers were incubated with rabbit polyclonal primary antibodies against occludin (BD Biosciences, 611091) or claudin-3 (Invitrogen, 34-1700), diluted 1:100 in PBS containing 5% goat serum and 0.1% Tween, for 1 h at room temperature. Monolayers were washed three times with PBS containing 0.1% Tween, with the final wash lasting 10 min, and incubated with Alexa Fluor 488-conjugated anti-rabbit secondary antibody (Invitrogen, A11034), diluted 1:100 in the same antibody diluent, for 30 min at room temperature protected from light.

Following the same washing procedure, insert membranes were excised and mounted cell-side up on glass slides using Fluoroshield with DAPI (F6057, Sigma) mounting medium. Coverslips were sealed with nail polish. Images were acquired using SP8 Leica confocal (Oil X63 objective, LasX acquisition software). Image processing and figure assembly were performed using Fiji (version 2.16.0).

## Statistics

TEER kinetics were analyzed using a two-way repeated-measures ANOVA with Geisser–Greenhouse correction, followed by Dunnett’s multiple comparisons test comparing each treatment condition with the untreated control at each time point.

## Supporting information

**Table S1: Bacterial strains and plasmids**

**Table S2: List of primers used in the study**

## Acknowledgments

We thank Frédérick Barreau’s team, particularly Sandrine Ménard, Amandine Roldan and Melvin Ayraud, for their contributions to the pilot experiments, and Laura Soler-Vasco and Olivier Puel’s team at Toxalim for providing access to equipment. We are grateful to all of them for their valuable discussions and advice throughout this work. This work was supported by the French National Agency for Research under grant VeSPath (ANR-23-CE14-0070-01) and AO Tremplin UT3 PIME-ID. Imaging experiments were performed at the Infinity-INSERM UMR1291 core facility connected to Toulouse Réseau Imagerie network, member of the France-BioImaging national infrastructure supported by the French National Research Agency (ANR-24-INBS-0005 FBI BIOGEN).

## Conflict of interests

The authors declare that they have no conflict of interest.

## Author contributions

**Conceptualization:** Marie Pénary, Eric Oswald, Priscilla Branchu

**Data curation:** NA

**Formal analysis:** Marie Pénary, Priscilla Branchu

**Funding acquisition:** Eric Oswald, Priscilla Branchu

**Investigation:** Marie Pénary, Christelle Marrauld, Priscilla Branchu

**Methodology:** Marie Pénary, Eric Oswald, Priscilla Branchu

**Project administration:** Priscilla Branchu

**Resources:** Marie Pénary

**Software:** NA

**Supervision:** Eric Oswald, Priscilla Branchu

**Validation:** Marie Pénary, Christelle Marrauld, Priscilla Branchu

**Visualization:** Priscilla Branchu

**Writing – original draft:** Priscilla Branchu

**Writing – review & editing:** Marie Pénary, Christelle Marrauld, Eric Oswald, Priscilla Branchu

## Notes

### Competing Interest Statement

The authors have declared no competing interest.

